# Rapid behavioural response of urban birds to COVID-19 lockdown

**DOI:** 10.1101/2020.09.25.313148

**Authors:** Oscar Gordo, Lluís Brotons, Sergi Herrando, Gabriel Gargallo

## Abstract

Biodiversity is threatened by the growth of urban areas. However, it is still poorly understood how animals can cope with and adapt to these rapid and dramatic transformations of natural environments. The COVID-19 pandemic provides us with a unique opportunity to unveil the mechanisms involved in this process. Lockdown measures imposed in most countries are causing an unprecedented reduction of human activities giving us an experimental setting to assess the effects of our lifestyle on biodiversity. We studied the birds’ response to the population lockdown by using more than 126,000 bird records collected by a citizen science project in north eastern Spain. We compared the occurrence and detectability of birds during the spring 2020 lockdown with baseline data from previous years in the same urban areas and dates. We found that birds did not increase their probability of occurrence in urban areas during the lockdown, refuting the hypothesis that nature has recovered its space in human emptied urban areas. However, we found an increase in bird detectability, especially during early morning, suggesting a rapid change in the birds’ daily routines in response to quieter and less crowded cities. In conclusion, urban birds showed high behavioural plasticity to rapidly adjust to novel environmental conditions, as those imposed by the COVID-19.

## 1. Introduction

Since the first human settlements some millennia ago, the anthropogenic transformation of the natural environment to build towns and cities has been a hallmark of humanity. During the last century, urbanization has experienced exponential growth across the world and it is expected to continue as more people will move from rural to urban areas [1-4]. As a result, urbanization has become one of the most important drivers of global change and a major threat to biodiversity [2,4-6]. Novel, human created environments, such as urban areas, represent a formidable challenge for organisms because the magnitude and peace of the environmental alterations imposed by humans usually exceed their limits of tolerance leading to populations shrinkage and extinction [5,7]. Urban challenges include dealing with chemical [3], acoustic [8,9] and light pollution [10,11], human disturbance [5,12], new pathogens [13,14] and predators [15,16], and human infrastructures [15,17]. However, some species are able to overcome these challenges and thrive in urban environments [4,7,12,18,19]. Therefore, a key question in urban ecology is how species cope with urbanization. Countless studies have demonstrated that adapting to urban environments imply some kind of phenotypical differentiation from non-urban relatives [7-9,12,18]. Indeed, organisms are forced to adjust their physiology, behaviour and life histories to the novel conditions imposed by the city [5,7]. However, little is known about the adaptive mechanisms allowing the differences observed between urban and non-urban dwellers [6,7]. Observed adjustments are mostly consistent with phenotypically plastic responses [12], but individual sorting and microevolutionary changes by divergent selection could be playing a role [4,5,7,18,19]. Perhaps, our inability to disentangle these mechanisms comes from a deficit of experimental studies in urban ecology [9], in spite of the fact that human transformed environments provide often ready-made experiments. The current spread of the novel coronavirus disease (COVID-19) and its consequences represents an excellent example, as we are involuntarily involved in a major unintended social experiment.

After the declaration of the COVID-19 pandemic in March 2020 by the World Health Organization, most countries have implemented social and health measures unprecedented in recent history. These measures, aimed at containing the virus spread [20-23], have focused on social distancing and population confinement, as well as the cease of non-essential productive and social activities. Overall, the measures have contributed to a global diminishing of human activities [24]. This abrupt and dramatic disruption of most human social and economic activities have already had quantifiable effects on urban environments by marked reductions in air pollution [25-27] and noise [28-30]. One of the most noticeable and generalized measure has been applying certain degree of population lockdown, which renders our city streets empty and virtually silent. This situation provides a once-in-a-lifetime opportunity to study urban wildlife responses to less crowded, noisy and polluted cities and gain unprecedented mechanistic insights into how human activities affect wildlife [24,31-33]. Product of the human lockdown, unusual observations of animals in urban areas worldwide have flooded the media and social networks planting in the social imaginary the idea that “nature is getting back its space” (*sensu* [34]). Although plausible, this idea is, in most cases, based on anecdotal records, sometimes false [34,35], without any quantitative scientific investigation supporting such claim [24,33].

In this work, we aimed to assess the behavioural responses of birds to the sudden and drastic changes occurring in urban environments resulting from the COVID-19 lockdown in a densely populated area of north eastern Spain (Catalonia). Following China [23] and Italy [21], Spain was the third country worldwide to impose a severe population lockdown to stop COVID-19 spread. The declaration of the national emergency in March 14^th^ 2020 by the Spanish Government imposed the strictest lockdown measures in Europe. Since then, social restrictions were alleviated progressively until the end of June (electronic supplementary material, figure S1, table S1). As in other parts of the world, this big halt of human activities has had significant environmental effects with reduced air contamination and noise in Spanish cities [26,27,36,37]. The severity of the lockdown measures imposed in Spain, make this country especially suitable to study COVID-19 lockdown effects in urban fauna, as they enjoyed exceptionally quieter and peaceful towns and cities during many weeks.

We compared bird records collected during the first four weeks of the lockdown in towns and cities of Catalonia (NE Spain) with the available records for the same region and dates since 2015. These historical records were used as baseline data. Our broad scale approach (hundreds of study sites covering and area of 32,000 km^2^) at community level (we studied 16 different species) allowed us a robust testing of two key questions:

1. Did urban birds become more common in response to human empty cities? It can be predicted that decreased human presence and disturbance allowed animals to occupy spaces that used to be above their fear tolerance thresholds [5,34,35]. Therefore, we expected a higher occurrence in 2020 compared to the historical records for the same urban areas. This effect being likely stronger for shier species (i.e., urban adapters), who are less tolerant to human disturbances [5,12,38].
2. Were urban birds more detectable as a consequence of quieter cities? It can be predicted that decreased anthropogenic noise increased the effective distance of among bird communications [5,8,9,12] and be more easily perceived by observers [39,40]. Moreover, as the masking effect of human acoustic contamination mostly disappeared, we expected an increase in singing activity, including potential shifts in its timing, to profit from the new urban soundscape [8,9,41,42]. Therefore, we expected a higher detectability of urban birds during the lockdown than in previous years, with possible changes in the daily patterns of detection.

## 2. Material and methods

### a) Bird data

On March14^th^ 2020, the Spanish Government declared the national emergency due to COVID-19 outbreak and imposed severe social restrictions. These restrictions included mandatory and permanent confinement of the population, borders closure, limitations in public transport, on-line education, working from home whenever possible, and closure of non-essential business and public services. One day later, we launched the project “#JoEmQuedoACasa” (I stay at home) within the citizen science on-line platform *ornitho* (www.ornitho.cat). This platform aims to collect wildlife records in Catalonia from birdwatchers and naturalists to improve knowledge of biodiversity in this region. *Ornitho* has been running since 2009 and has gathered more than 6.5 million records to date. The project launched during the lockdown aimed to collect information about wildlife responses to the new environmental conditions resulting from people confinement. In addition to this valuable information, the project was important to keep engaged birdwatchers in this citizen science program by encouraging them to continue complete checklists submission, even during a period of constrained outdoor activities [43-45]. A complete checklist is a checklist with all identified species during any survey.

Lockdown surveys were conducted between March 15^th^ and April 13^th^ of 2020. During these four weeks, people was subjected to the most restrictive conditions of mobility and consequently this period showed the most drastic reduction of human activities (electronic supplementary material, figure S1, table S1). Therefore, lockdown checklists were carried out only from homes (e.g., balconies, rooftops or yards). To determine the effect of lockdown on bird behaviour, we also gathered all complete checklists available in *ornitho* recorded during the same dates between 2015 and 2019. Surveyed sites were classified as urban or non-urban according to the 2017 land use/land cover map of Catalonia [46]. All surveys during the lockdown were in urban environments, except a few observers living in the countryside, which were excluded from the analyses. Therefore, we obtained three groups of checklists: urban lockdown, historical urban, and historical non-urban, which contained a total of 126,315 bird records. Historical urban data represented baseline data, while historical non-urban data was included as control data without human disturbances. We used five years of historical data together to have a comparable number of checklists in urban areas to those recorded during the lockdown. By using several years of historical data we got a more representative baseline of the usual conditions previous to the COVID-19 pandemics, although we could not assess variability among years.

All checklists had associated basic information about the survey: site (geographical coordinates), date, hour, time invested (which was used as a proxy for sampling effort) and observer identity. We excluded checklists lasting >3 h, as they might be discontinuous surveys. We also excluded those checklists started one hour earlier or later than sunrise or sunset, respectively, as they represented nocturnal surveys. To correct for the adjustment of daylight saving time at the end of March, we rescaled recorded hours in civil time to the relevant daily sun events: sunrise, noon and sunset, which were established as -1, 0 and 1, respectively. Sunrise, noon and sunset were calculated for every geographical coordinate and date by the ‘suncalc’ library (version 0.5.0) for R software [47]. Rescaling was calculated as the quotient between the difference of noon and checklist hour and the difference of sunrise or sunset and checklist hour, depending on whether checklist started earlier or later than noon, respectively. This transformation allowed to fix the small bias caused by the longitudinal differences in sunrise and sunset across Catalonia as well as by the progressive day length increase during the study period. Not many observers recorded the number of individuals for each species. For this reason, we opted to work with presence/absence data.

We gathered 1,289 complete bird checklists at 149 sites for the lockdown period. The number of replicated surveys per site and observer ranged from 1 to 91 (mean=8.7, SD=12.4). Historical records in urban areas were the scarcest: 1,062 checklists in 410 sites with up to 48 replicates per site (mean=2.6, SD=5.2). As expected, data from non-urban areas were the most abundant, as observers usually preferred birdwatching in natural habitats. We gathered 5,849 checklists from 3,113 sites. Although one observer made 84 replicates for the same site, on average, observers in this group showed the lowest site fidelity (mean replicates=1.9, SD=3.6).

We selected data for the 16 most common sedentary urban species in Catalonia [48,49] (see figure 1). We focused only on sedentary birds to avoid seasonal changes in occurrence and abundance associated with migration. Data from the common and the spotless starlings (*Sturnus vulgaris* and *S. unicolor*, respectively) were merged as *Sturnus* spp. as both were not usually identified at species level in most observations due to their high resemblance [50]. Both species are common, well spread, sympatric and share similar habits and behaviour [49]. Thus, we did not expect important differences in their occurrence or detectability.

**Figure 1.**
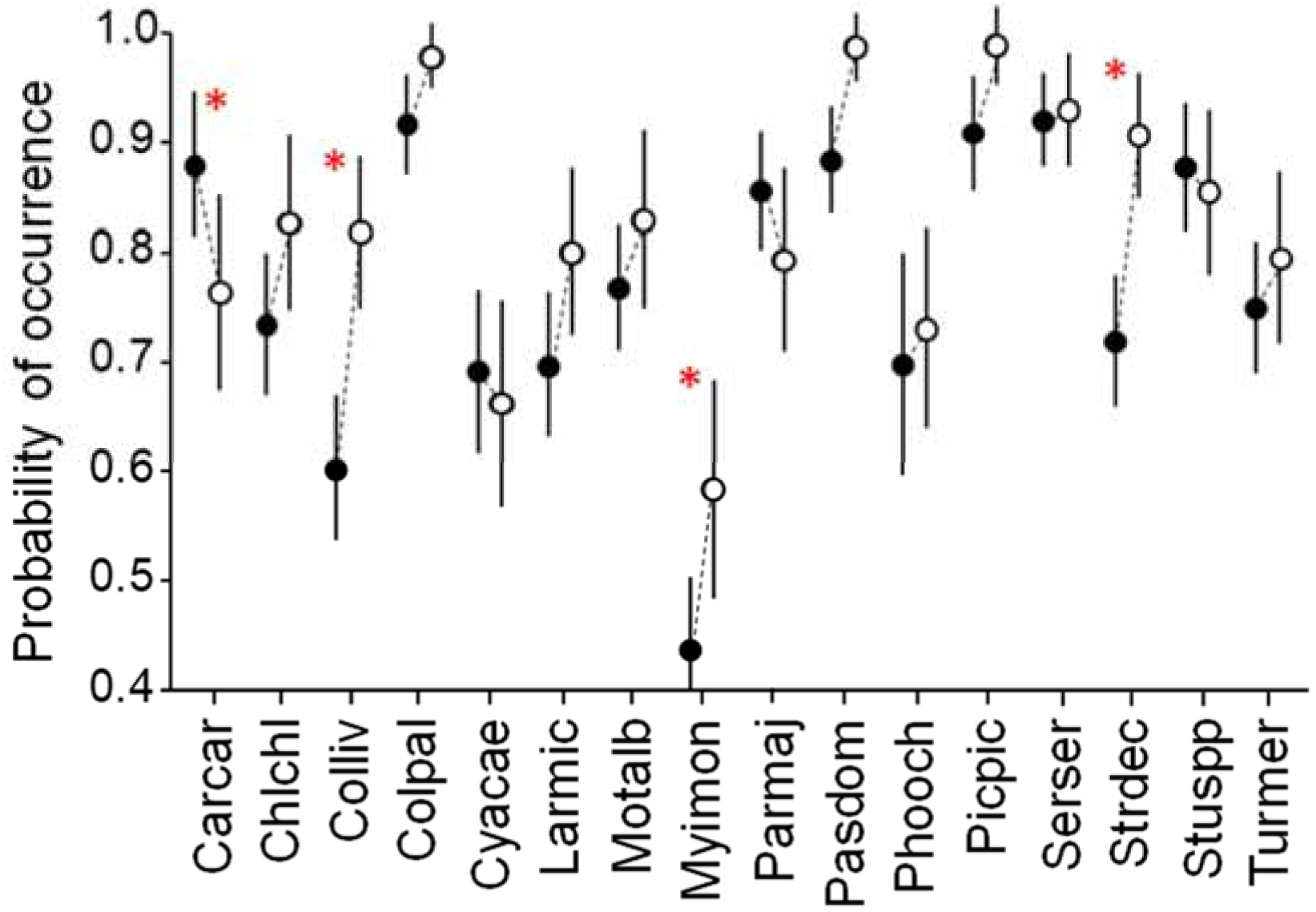
Probability of occurrence of birds in urban areas before (2015-19, black dots) and during (2020; white dots) the COVID-19 lockdown. Asterisks indicate significant differences (*p*-value < 0.05). Error bars denote 95% confidence intervals. Acronyms for the species: Carcar *Carduelis carduelis*, Chlchl *Chloris chloris*, Colliv *Columba livia*, Colpal *Columba palumbus*, Cyacae *Cyanistes caeruleus*, Larmic *Larus michahellis*, Motalb *Motacilla alba*, Myimon *Myiopsitta monachus*, Parmaj *Parus major*, Pasdom *Passer domesticus*, Phooch *Phoenicurus ochruros*, Picpic *Pica pica*, Serser *Serinus serinus*, Strdec *Streptopelia decaocto*, Stuspp *Sturnus* spp., Turmer *Turdus merula*.

### b) Statistical analyses

To disentangle the effects of individuals’ presence (first question) and detection (second question) in our bird data, we used hierarchical occupancy models [51,52]. We considered as replicated surveys those checklists reported by the same observer within the same 1×1 km UTM cell. By combining observer and location, we avoided variability in detection rates due to observer expertise. We could assume confidently that observer experience was randomly distributed across our study area. The equations defining our model were:

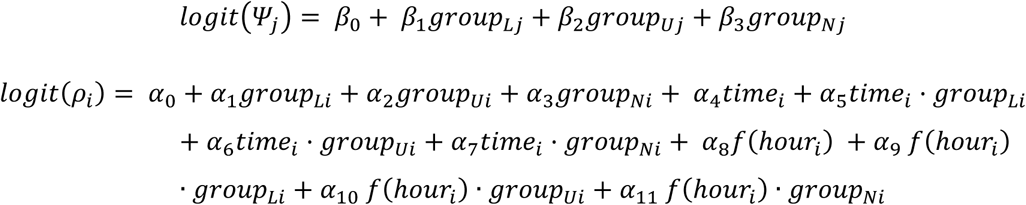

where *Ψ*_*j*_ is the occurrence of a species at site *j* and *<u>ρ</u>*_*i*_ is its detectability in the checklist *i*; groups *L, U* and *N* refer to lockdown, urban historical and non-urban, respectively; time refers to the duration of the survey; and hour refers to the starting hour. Hour was included as an unpenalized thin plate regression spline basis function (*f*) with five degrees of freedom because we expected that detectability could vary in a non-lineal way along the day [53,54]. Interactions between group and time and between group and hour allowed to model the effect of these two variables on detectability within each group. To test the significance of hour and interactions, we used log likelihood ratio tests. Basis functions were built by the smooth.construct function from package ‘mgcv’ (version 1.8-22 [55]), while occupancy models were run with the occu function of package ‘unmarked’ (version 0.12-3 [56]) for R.

## 3. Results

Probability of occurrence of a species during the lockdown did not differ significantly from the occurrence recorded in urban areas in previous years in 12 out of the 16 studied species after accounting for their imperfect detection (figure 1; electronic supplementary material, table S2). In the four species with significant differences, three increased their occurrence and one decreased it. As expected, most of the species (10) showed significant differences in their occurrence between lockdown and non-urban checklists (electronic supplementary material, table S2). On average, these species were approximately a 15% more common in the lockdown checklists than in the non-urban checklists, confirming that most of the studied species were preferentially urban dwellers.

For most species (10), probability of detection was higher in lockdown checklists than in historical urban ones, but this difference was not statistically significant in most cases (electronic supplementary material, figure S2, table S3). Most species were less detectable in non-urban checklists than in urban ones.

As we predicted, detectability varied along the day in a non-linear way for all species (figure 2; electronic supplementary material, figure S3). Except for two species, the pattern of daily variation in detectability was significantly different among groups (electronic supplementary material, table S4). A difference consistently found in most species was higher detectability in the first hours of the morning during the lockdown compared to the urban records from previous years (figure 2, 3). In most species, during the lockdown detectability peaked at dawn and decreased until midday, while in the historical urban checklists the peak of detectability was around mid-morning. In fact, the pattern of detectability along the day in the lockdown group resembled more to the non-urban pattern than to the urban pattern in many species. Predicted detectability at sunrise by our models in the lockdown group was on average a 27% higher than in the urban group (sign test: *Z*=3.25, *p*-value=0.001; figure 3).

**Figure 2.**
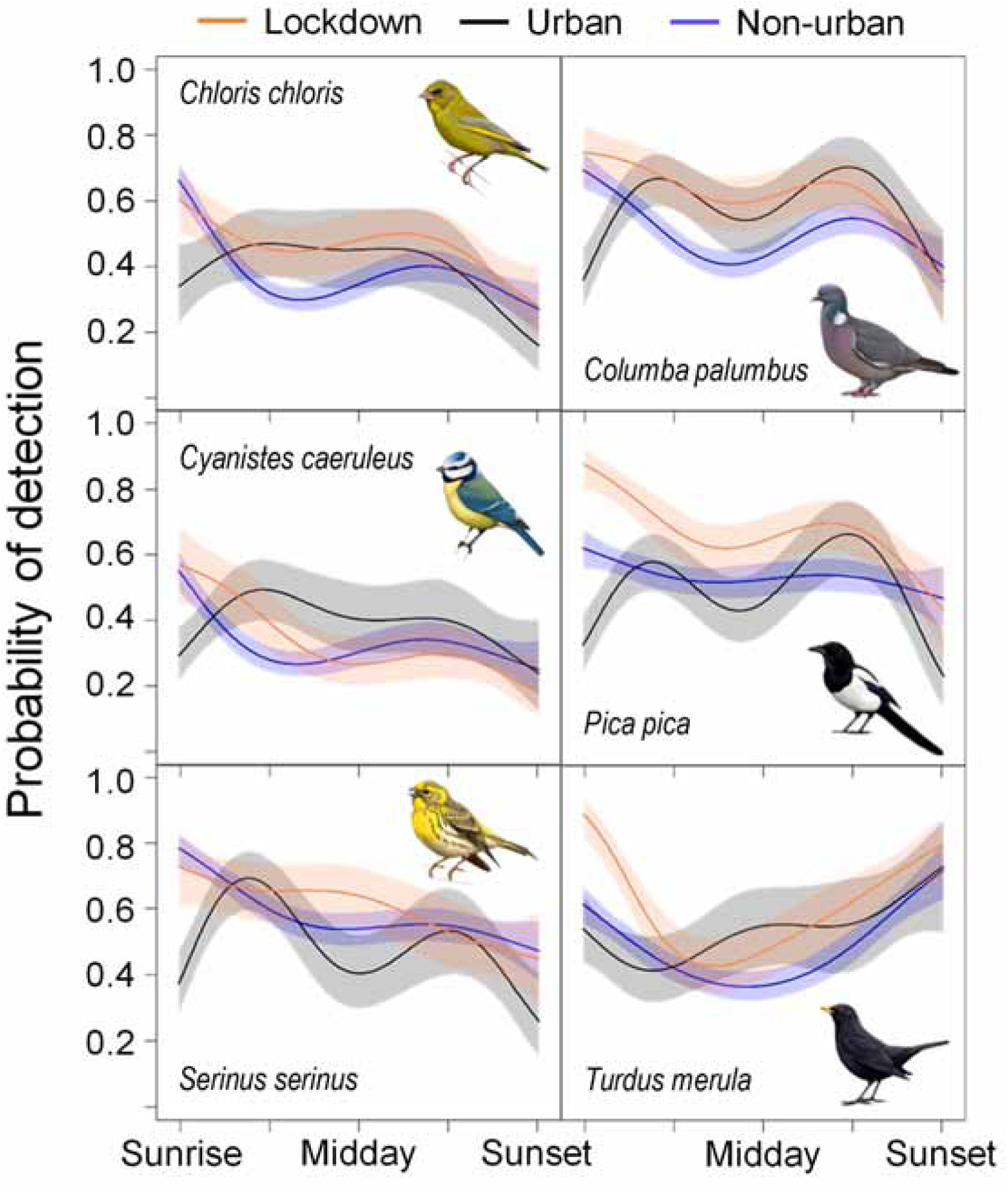
Variation in the probability of detection along de day for each group of data (collected during the lockdown, collected historically in urban sites, and collected in non-urban environments). Shaded areas represent the 95% confidence intervals. See fig. S2 in the electronic supplementary material for the rest of species. Bird illustrations by Martí Franch/Catalan Ornithological Institute.

**Figure 3.**
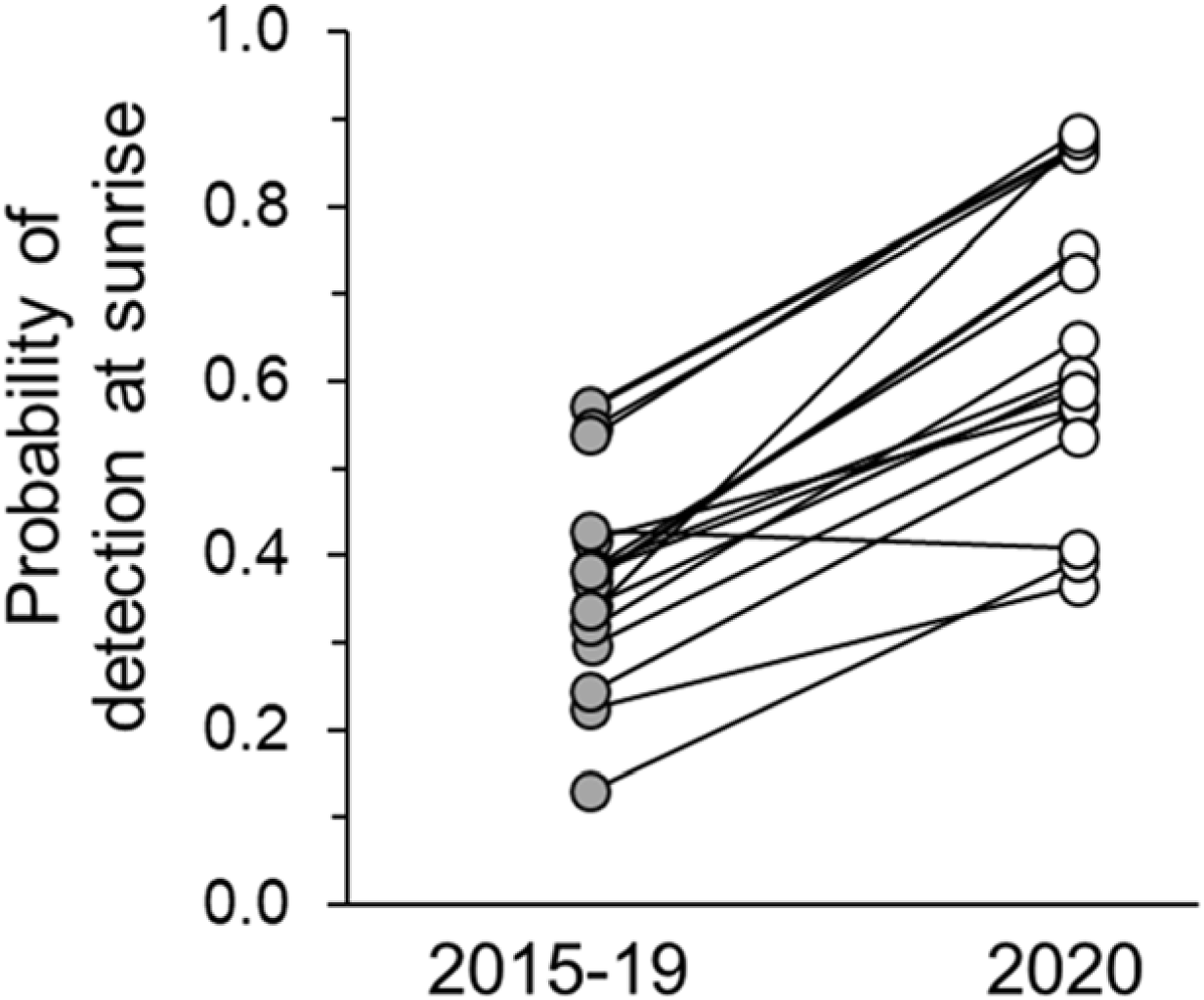
Probability of detection in urban environments at sunrise for the 16 studied bird species before (2015-19) and during (2020) the COVID-19 lockdown.

As expected, in all but one species, chances of detection increased with longer surveys (electronic supplementary material, figure S4, table S5). In most of them (11), such time effect was significantly different among groups (electronic supplementary material, table S4). Therefore, a certain increase of the sampling time implied a different increase in chances of detection in the lockdown, the historical urban and the historical non-urban groups for most species. For half of the species, sampling time effect was significantly lower in the lockdown group than in the historical urban group (mean reduction of 17%; electronic supplementary material, table S5). This systematic reduction contrasts with the comparison of time effect between lockdown and non-urban groups, where for nine species there were significant differences between both groups, but such differences were disparate (mean change -0.8%; electronic supplementary material, table S5).

## Discussion

Birds did not occur in higher rates in towns and cities during the lockdown than before it, not supporting the hypothesis that birds moved into the human emptied urban areas [33,34,38]. As the changes induced by the COVID-19 lockdown were drastic and sudden and did not last enough, they probably did not allow for colonization processes. The few species with a significant increase of their prevalence in urban surveys during lockdown were, interestingly, the ones that are mostly urban. As these species are not present in large numbers away from urban areas, they could hardly rely on non-urban source populations to occupy cities and towns during the lockdown. Their occurrence in a higher proportion of checklists during the lockdown could be due to the observers being constrained to survey from their homes. Urban checklists recorded during the lockdown probably were more focused on more extreme urban environments (i.e. core urban areas) than those checklists historically recorded in urban areas of Catalonia, which perhaps included a greater proportion of surveys in urban parks, landscaped plots or suburbs. These areas are considered urban areas from a land-use perspective, but they have more diversity of habitats at our working micro-scale (1×1 km), where the most urban exploiter species, such as feral pigeons (*Columba livia*), collared doves (*Streptopelia decaocto*) and monk parakeets (*Myiopsitta monachus*), may not find their most suitable niche. The absence of some non-essential activities during the lockdown, such as feral populations’ management and culling [32], can be discarded as a cause for the increase of occurrence of these species. Despite being harmful invasive species [57,58], there is no management of the Catalan populations of Psittacidae yet. Culling of feral pigeons in big cities, as Barcelona, was suspended due to ethical considerations in 2006 [59] and thus, control measures for this species were unaffected by the lockdown. Finally, the collared dove colonized naturally Catalonia in the 70s and occupied quickly most of the urban areas during the 80s and 90s [49,60]. Currently, its population grows at a 2% annual rate [61], which is too small to justify a 18% increase of population occupancy in 2020.

Birds changed their detectability pattern throughout the day as a consequence of the lockdown. In general, there was an increase in detection probability, which was especially marked in the early morning. As observed in non-urban habitats, detectability during the lockdown decreased from dawn onwards, while at the same urban locations detectability was historically low at dawn and increased until reaching a peak two or three hours later. It is interesting to note that the Eurasian blackbird, a model species in urban ecology studies [7,8,12,62,63], was the only exception to this pattern. Overall, many species showed a pattern of detection during the lockdown in urban areas more similar to that commonly seen on natural environments. Although in urban areas the 2020 detectability pattern was patently different from the 2015-2019 baseline level, such differences could have been partially enhanced because of comparing a single year against several pooled years. Nevertheless, it would be necessary to know whether or not there is significant among years’ variability in bird daily routines. If that is the case, we would expect that behavioural responses to extreme events, such as the covid-19 lockdown, would fall out of the normal variability. Unfortunately, we could not properly explore the between years’ variation in detectability patterns due to sample size constrains.

Urban birds during lockdown may have shown this detectability peak at dawn, typical of non-urban habitats, because of a rapid behavioural response to adjust to the new environmental conditions imposed by the COVID-19 measures [64-68]. Birds rely heavily on acoustic communication. During reproduction, males sign to attract females and defend their territories, becoming highly conspicuous and detectable. COVID-19 lockdown was imposed just at the beginning of the breeding season, when singing activity was expected to be especially high [69]. Therefore, there was a strong pressure to time singing activity to the optimal moment of the day. This moment is dawn because the physical properties of the atmosphere enhance acoustic transmission [70,71] and consequently, birds can reach the maximum audience. Thus, urban birds during the lockdown may have advanced their main period of singing activity to dawn, increasing their detection at those hours, similar to what is observed in non-urban areas.

During the lockdown, human presence and activities decreased drastically (electronic supplementary material, figure S1, table S1), being this especially notable during rush hours, which almost disappeared [27,30,36]. During the spring in Spain, morning rush hour matches with the first hours of light, when birds are expected to be especially communicative [42,63,72]. The dramatic decrease in noise during the lockdown released early morning acoustic space that could be recovered by the dawn chorus. Empirical and experimental evidence demonstrates that urban birds avoid the masking effect of anthropogenic noise [8,9,41,42,73]. Our findings match these previous studies, but instead of advancing the dawn chorus [42,62,63,74], our historical urban data suggests that birds would delay their peak of activity (and consequently of detectability) to mid-morning. In our study context, this can be explained because civil and solar time are heavily decoupled in Spain since the country is located in the westernmost part of its time zone [75]. For this reason, if birds in Catalonia advance their activities before sunrise, they would be still suffering an important overlap with morning noisy human activities, such as commuting, school attendance, shop opening, etc. [63,72]. Hence, the best option for birds would be to delay the peak of activity to after the morning rush hour [73]. Moreover, most of the previous studies have been carried out in more northern latitudes [8,62,76], where climate conditions can still be severe at night in early spring. Under these circumstances, individual survival can be challenged by a strong nocturnal energy demand [77,78]. There, dawn singing can become a relevant and honest signal of phenotypic quality of males, as only those individuals in best physical condition can undergo dawn fasting [62]. In Mediterranean regions, where spring nights are mild, the role of dawn singing as signal of male quality might be less important. Attracting mates would be the prime objective for singing and consequently, males would be more pressed to place this activity when interference of anthropogenic noise is at its lowest. Since sunrise, these lowest levels of noise are just after the morning rush hours (i.e. later than 9 a.m.), when the air physical properties still keep sound attenuation and fluctuation low [70].

If birds have changed their behaviour, this adaptive, flexible behavioural response must have been mediated by phenotypic plasticity. People lockdown was sudden and the environmental scenario in urban areas changed radically from one day to the next (electronic supplementary material, figure S1) [27,36,37]. This unprecedented social experiment imposed by the COVID-19 allowed us to test and support the hypothesis of the high plasticity displayed by individuals living in urban areas in order to cope with a constantly changing environment [5-7,64]. However, this fast adaptive response might have been facilitated by a previous conditioning of birds to weekly rhythms of human activities. Birds change their behaviour from working to weekend days to match with human behaviours [41,73,79]. Therefore, birds could assimilate the lockdown as a very long and especially peaceful weekend and consequently, we may speculate that behavioural adjustments to novel lockdown conditions happened quickly, necessarily in less than one month. Nevertheless, it would be interesting to explore the long-lasting consequence at a community level of this environmental change [76]. Weekends are just two days long, while strict people lockdown lasted for at least two months in most regions of Spain. One may speculate that bolder and fast-adapting species are able to modify their behaviour on a weekly basis. However, during the lockdown, all species had enough time to habituate to the long-lasting new conditions. In fact, as we have demonstrated, all of them modified their daily patterns of detectability. Maybe the most urbanite species have benefited the least of this lockdown as their boldness and higher human tolerance was no longer an advantage in empty cities.

In addition to the birds’ rapid behavioural response to the anomalous environmental conditions during the lockdown, observers had certainly enhanced opportunities to detect birds during this period. Urban areas were quieter than usual [27-30], improving the chances of listening the birds [34,39,40,70]. Moreover, absence of people outdoors allowed for the display of shy and distrustful behaviours [5], facilitating bird observations, especially for those less singing species, as the magpie (*Pica pica*) or the yellow-legged gull (*Larus michahellis*). Hence, birds’ detectability during 2020 could be higher just as a by-product of a reduced interference in urban birdwatching of the human activities during the shutdown (e.g., traffic, pedestrians, factories, etc. [30,36,37]). However, these improved conditions for urban birdwatching were heavily constrained by the fact that observers were forced to stay at their homes and their sampling area was reduced to what they could see from there. Therefore, improved detection was to some extent counterbalanced by the limited scope from the survey sites. The observed effect in increased sampling time would support this hypothesis, as we demonstrated that the discovery rate in most species was slower during the lockdown than in the historical urban surveys.

The differences observed between urban and non-urban environments were expected as habitat configuration and bird densities are patently different between them. In fact, populations of urban exploiter birds show usually higher densities in cities than in rural or natural close areas [5-7], facilitating their detection in urban areas. Such differences may have serious consequences for monitoring schemes aiming to quantify wildlife occurrence and abundance by standardized protocols, as the assumption of equal detectability under similar circumstances is usually violated [39,52,53,80]. For instance, one sampling hour at dawn is not equivalent in terms of chances to detect a species in urban and non-urban habitats. Traditional protocols assume that the best moment to detect birds is early morning [70,81], which is actually true, but apparently only in natural conditions without human disturbance, as we have demonstrated here (see figure 2). If the detectability peak in most urban populations is reached at mid-morning, their abundance would be systematically underestimated by usual sampling protocols based on early morning bird surveys. No matters whether this lower detectability in urban areas in early morning is caused by different daily routines of urban vs non-urban populations or by the masking effect of human activities in the city (or a combination of both). As there is an increased awareness about the importance of urban populations for bird conservation [7,48], it is necessary to ensure its accurate quantification, which may imply a redefinition of the most popular current census techniques [7]. Additionally, in this work we demonstrated the utility of occupancy models and the necessity to account for imperfect detection [53,54].

The COVID-19 shutdown has revealed the stress, noise and pollution present in urban areas [25,26,28,29]. Under typical urban conditions, bird behaviour is apparently altered by our lifestyle and thus, the possibility to enjoy the natural values of our cities is notably diminished [7]. Our society should reflect on our urban lifestyle and how it affects welfare of urban fauna and jeopardizes its conservation. As the world is becoming more urbanized and animals will be forced to live more often in anthropogenic environments [5-7], one way to ensure their adaptation as urban dwellers would be by reducing our noisier and more disturbing activities. Most importantly, not only urban populations of non-human animals would be benefited, but also ourselves from quieter, more peaceful and less polluted cities.

## Acknowledgments

We warmly thanks to the thousands of observers who shared their observations by *ornitho* website, for their long-term involvement in this project, and the special efforts undertaken during the lockdown. *Ornitho* project is supported by the Generalitat de Catalunya. Margarida Barceló-Serra provided valuable English edits and suggestions. We would also like to thank the anonymous reviewers and the editor for their comments.

## ELECTRONIC SUPPLEMENTARY INFORMATION

**Figure S1**. Variation in people’s visiting habits to six categories of places in Catalonia between March 1^st^ and July1^st^.

**Figure S2**. Probability of detection of birds in urban areas before and during the COVID-19 lockdown

**Figure S3**. Variation in the probability of detection along the day for each group of data in 10 studied species.

**Figure S4**. Effect of sampling time on the probability of detection for each group of data.

**Figure S1.**
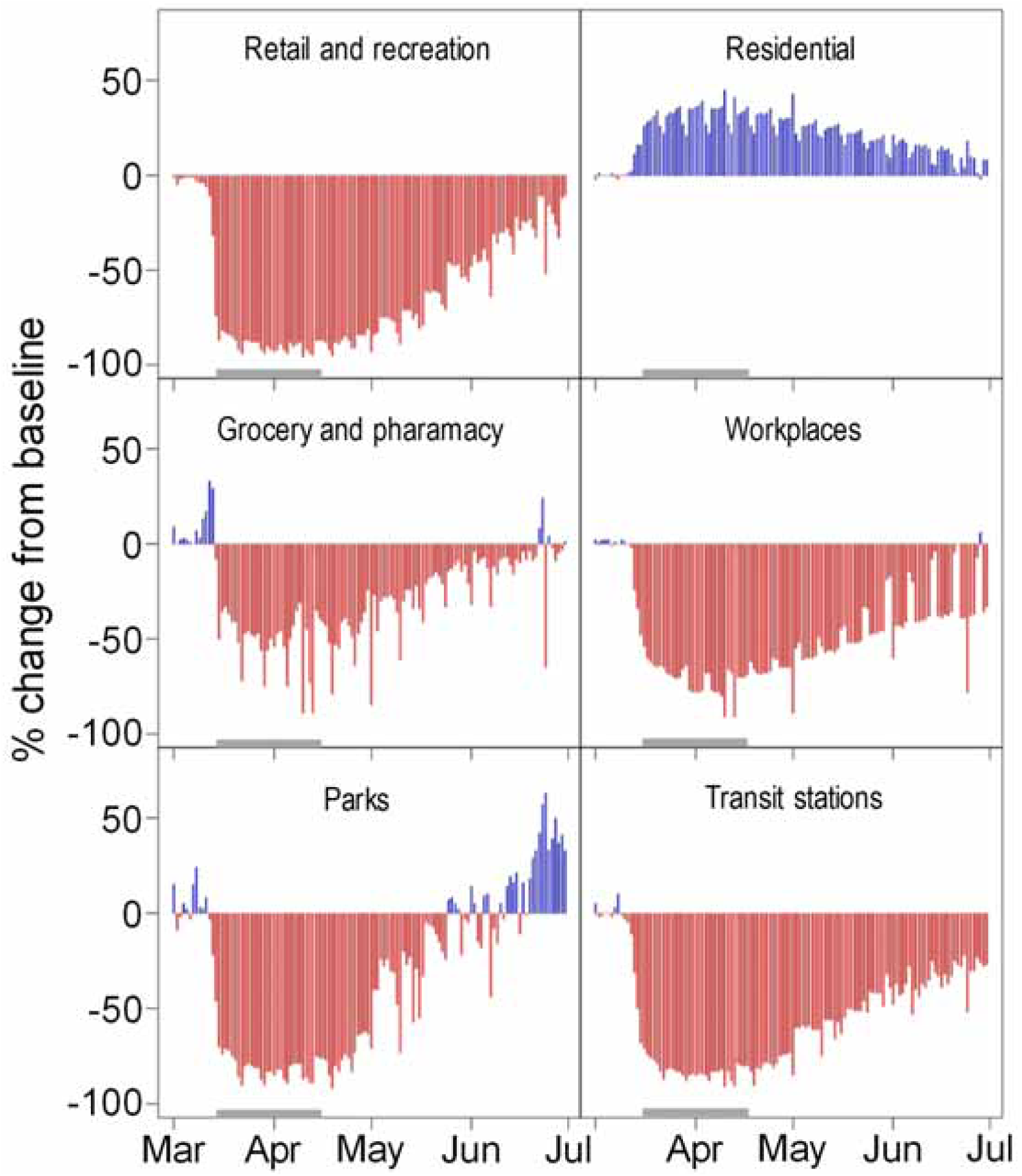
Variation in people’s visiting habits to six categories of places in Catalonia between March 1^st^ and July1^st^. These graphs can be interpreted as a proxy for human activity. To improve visualization, negative deviations are shown in red and positive in blue. The grey band on to the x axis shows the period of bird data collection. The emergency state in Spain due to COVID-19 lasted from March 15^th^ to June 21^st^. Note the forced strict lockdown during the first two months of this period and the following progressive recovery of activities. Figure data are freely available from “*Google COVID-19 Community Mobility Reports*” (https://www.google.com/covid19/mobility/ Accessed: August 25^th^ 2020). Baseline level (i.e., 0%) has been calculated as the average from January 3^rd^ to February 6^th^ 2020 (visit the previous link for further technical details on data calculations).

**Figure S2.**
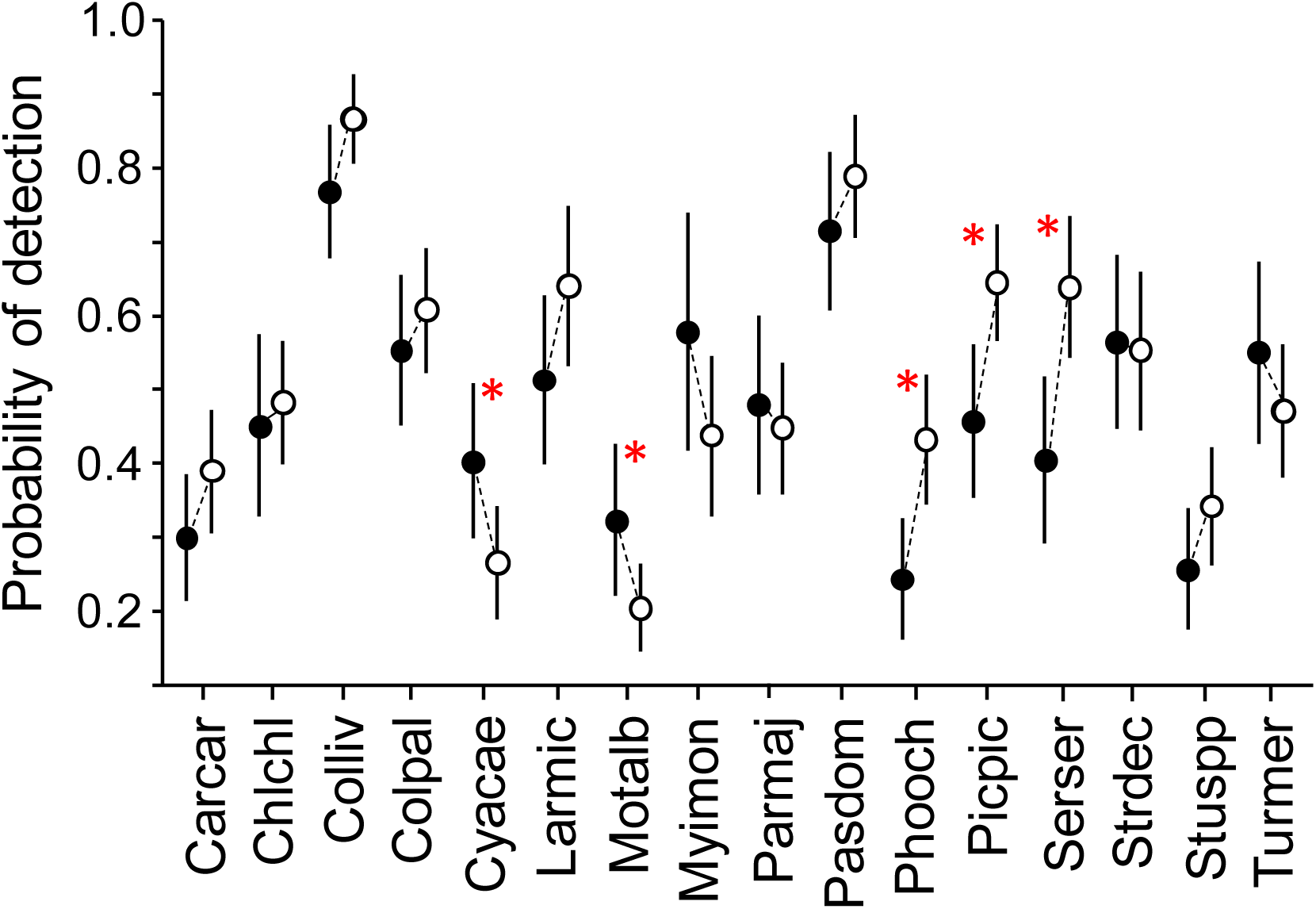
Probability of detection of birds in urban areas before (2015-19, black dots) and during (2020; white dots) the COVID-19 lockdown. Asterisks indicate significant differences (*p*-value < 0.05). Error bars denote 95% confidence intervals. Acronyms for the species: Carcar *Carduelis carduelis*, Chlchl *Chloris chloris*, Colliv *Columba livia*, Colpal *Columba palumbus*, Cyacae *Cyanistes caeruleus*, Larmic *Larus michahellis*, Motalb *Motacilla alba*, Myimon *Myiopsitta monachus*, Parmaj *Parus major*, Pasdom *Passer domesticus*, Phooch *Phoenicurus ochruros*, Picpic *Pica pica*, Serser *Serinus serinus*, Strdec *Streptopelia decaocto*, Stuspp *Sturnus* spp., Turmer *Turdus merula*.

**Figure S3.**
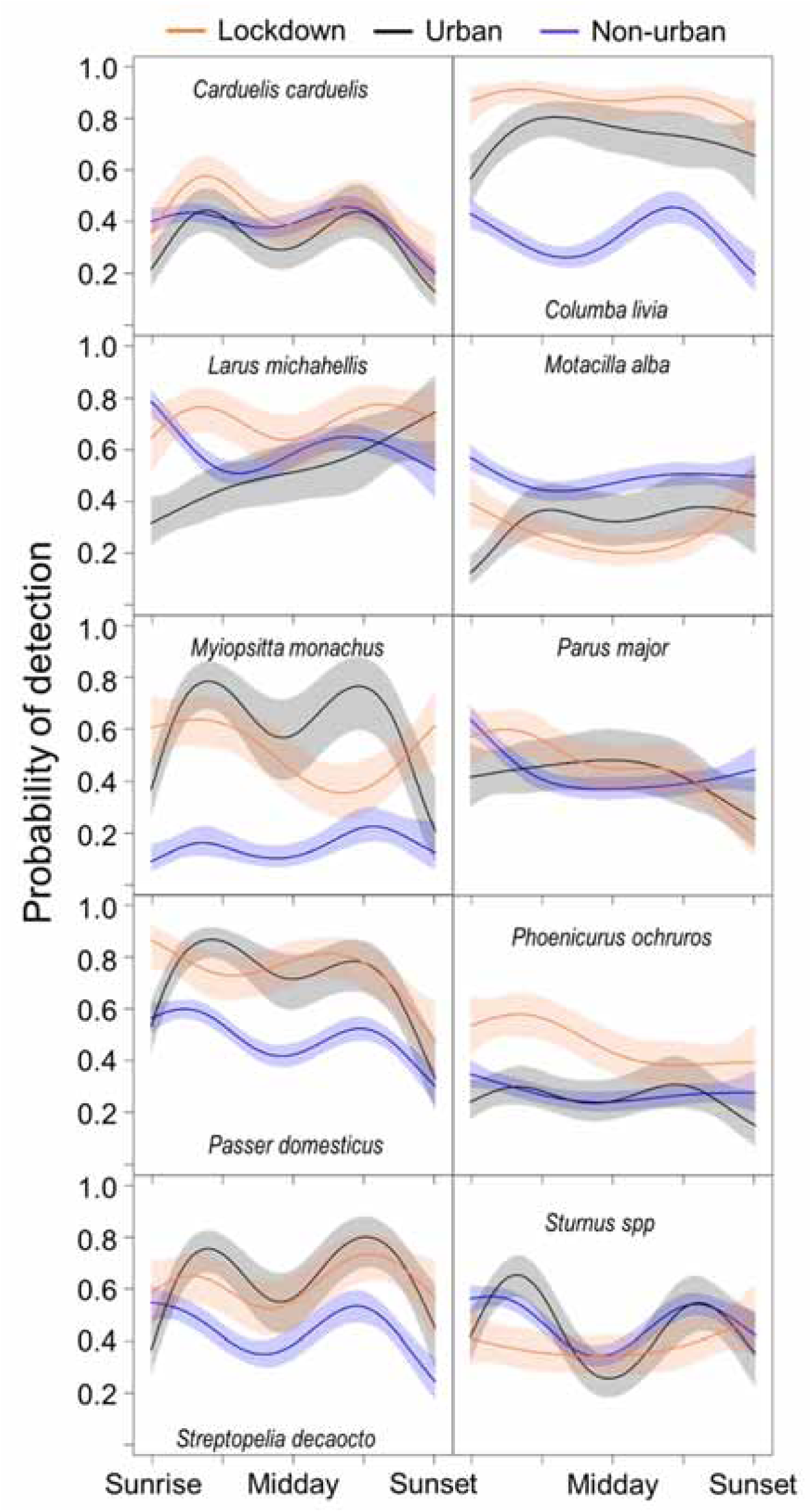
Variation in the probability of detection along de day for each group of data (collected during the lockdown, collected historically in urban sites, and collected in non-urban environments). Shaded areas represent the 95% confidence intervals.

**Figure S4.**
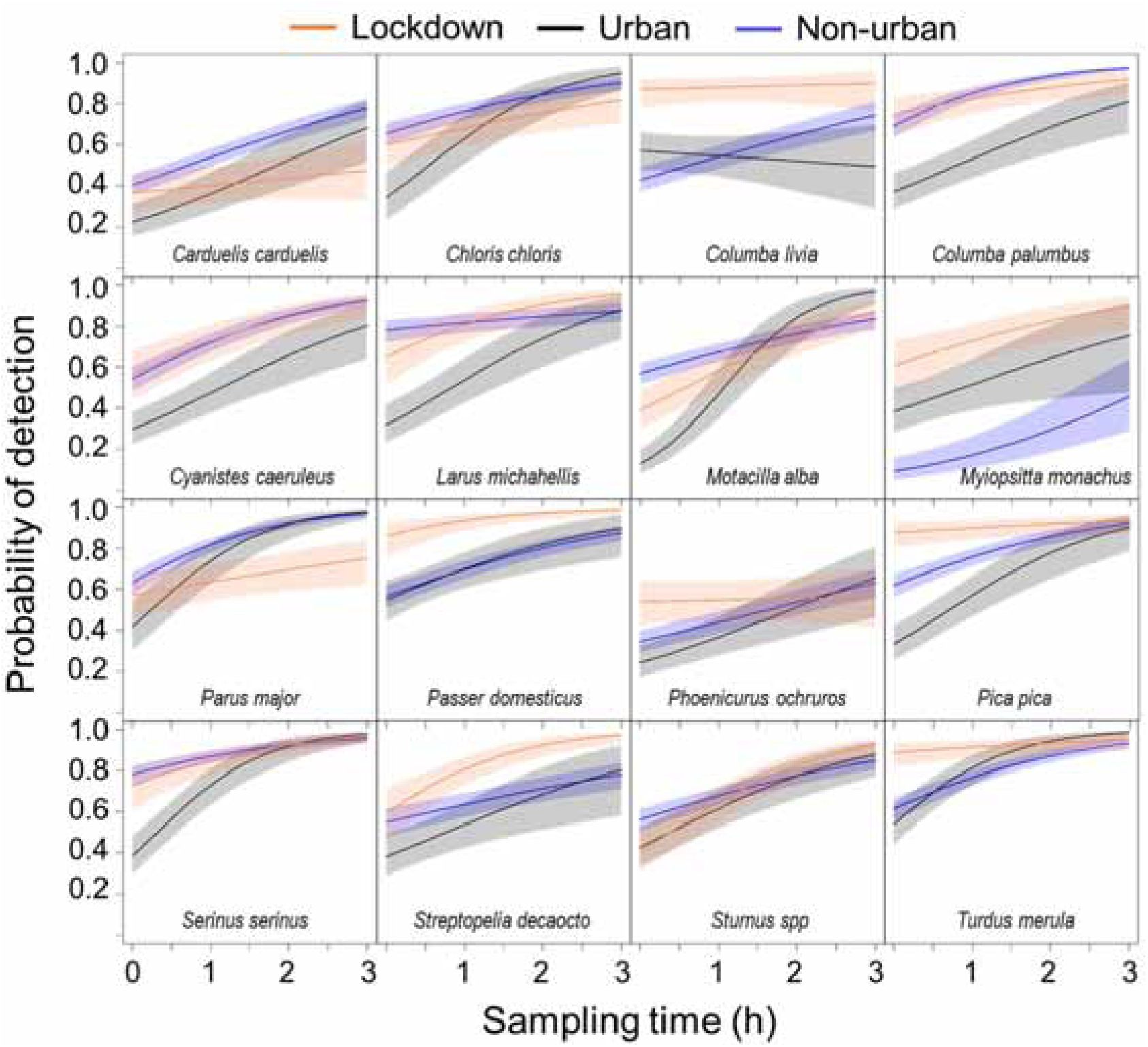
Effect of sampling time on the probability of detection for each group of data (collected during the lockdown, collected historically in urban sites, and collected in non-urban environments). Predictions done for surveys started at dawn. Shaded areas represent the 95% confidence intervals.

**Table S1.** Percentage variation in people’s visiting habits to six categories of places between March 15^th^ and April 14^th^ in Catalonia and its provinces. The values show the average of daily deviations from the baseline level. Outdoor sites experienced a severe decline, while people stayed more at home. See figure S1 for details.

| **Place** | **Catalonia** | **Barcelona** | **Girona** | **Lleida** | **Tarragona** |
| --- | --- | --- | --- | --- | --- |
| Retail and recreation | -89.56 | -90.41 | -88.13 | -86.19 | -88.56 |
| Grocery and pharmacy | -51.28 | -53.13 | -51.59 | -52.72 | -53.06 |
| Parks | -81.34 | -83.59 | -74.22 | -62.63 | -76.63 |
| Transit Stations | -82.41 | -82.44 | -80.41 | -78.88 | -83.00 |
| Workplaces | -70.41 | -72.22 | -64.22 | -61.81 | -65.69 |
| Residential | 31.25 | 32.03 | 28.41 | 26.00 | 27.84 |

**Table S2.**
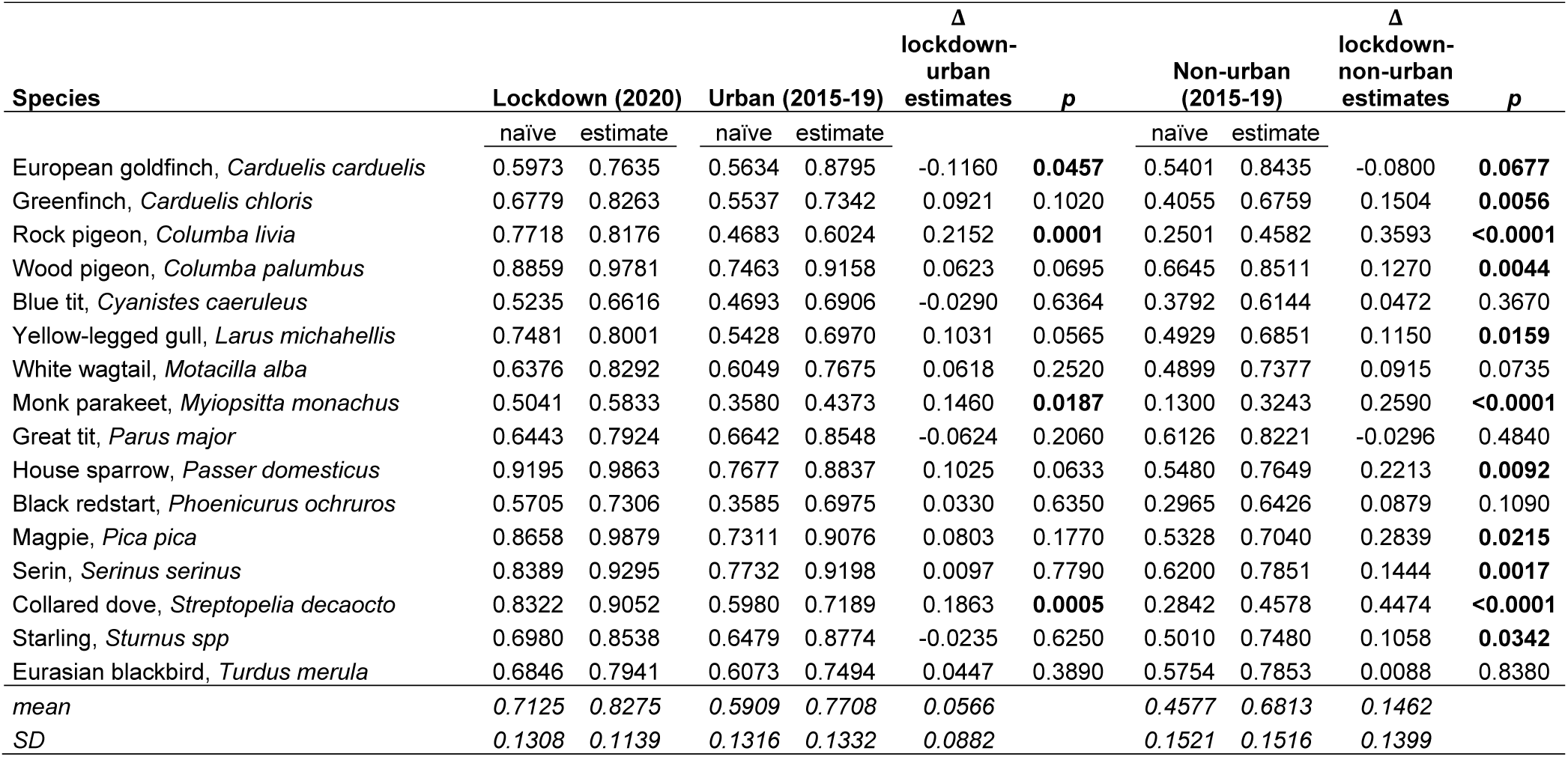
Results of the occupancy part of the occupancy models. *Naïve* columns show the proportion of sites where the species was found during the surveys, i.e. the raw occurrence without correction for the imperfect detection of birds. *Estimate* columns show the estimated occurrence of the studied species once the detectability was accounted for. As expected, estimated occupancy was always higher than the observed (i.e., *naïve*), demonstrating both imperfect and variable detection of birds. For the urban and non-urban groups of historical data (2015-19), a column with the differences (Δ) between their estimates and the lockdown group (2020 data) estimates is provided. The *p*-values for these differences are also shown. Values <0.05 are in bold. At the bottom of the table, the average and standard deviation of the studied species are given.

**Table S3.** Results for group effects on the detection part of the occupancy models. For the urban and non-urban groups, a column with the differences (Δ) between their estimates and the lockdown group estimates is provided. The *p*-values for these differences are also shown. Values <0.05 are in bold. At the bottom of the table, the average and standard deviation of the studied species are given.

| Species | Lockdown<br>(2020) | Urban<br>(2015-19) | $\Delta$ lockdown-<br>urban | $p$ | Non-urban<br>(2015-19) | $\Delta$ lockdown-<br>non-urban | $p$ |
| --- | --- | --- | --- | --- | --- | --- | --- |
| European goldfinch, *Carduelis carduelis* | 0.3897 | 0.3002 | 0.0895 | 0.1444 | 0.3948 | -0.0051 | 0.9133 |
| Greenfinch, *Carduelis chloris* | 0.4820 | 0.4517 | 0.0303 | 0.6938 | 0.3469 | 0.1351 | **0.0049** |
| Rock pigeon, *Columba livia* | 0.8665 | 0.7688 | 0.0978 | 0.0716 | 0.3289 | 0.5376 | **<0.0001** |
| Wood pigeon, *Columba palumbus* | 0.6070 | 0.5535 | 0.0534 | 0.4289 | 0.4302 | 0.1768 | **0.0005** |
| Blue tit, *Cyanistes caeruleus* | 0.2658 | 0.4035 | -0.1377 | **0.0364** | 0.3036 | -0.0378 | 0.4140 |
| Yellow-legged gull, *Larus michahellis* | 0.6405 | 0.5131 | 0.1274 | 0.1182 | 0.5865 | 0.0540 | 0.3912 |
| White wagtail, *Motacilla alba* | 0.2052 | 0.3233 | -0.1181 | **0.0419** | 0.4716 | -0.2664 | **<0.0001** |
| Monk parakeet, *Myiopsitta monachus* | 0.4378 | 0.5773 | -0.1395 | 0.1640 | 0.1097 | 0.3281 | **<0.0001** |
| Great tit, *Parus major* | 0.4479 | 0.4803 | -0.0324 | 0.6730 | 0.3717 | 0.0762 | 0.1290 |
| House sparrow, *Passer domesticus* | 0.7891 | 0.7150 | 0.0740 | 0.2820 | 0.4224 | 0.3667 | **<0.0001** |
| Black redstart, *Phoenicurus ochruros* | 0.4321 | 0.2437 | 0.1885 | **0.0028** | 0.2414 | 0.1907 | **<0.0001** |
| Magpie, *Pica pica* | 0.6452 | 0.4573 | 0.1879 | **0.0058** | 0.5259 | 0.1192 | **0.0169** |
| Serin, *Serinus serinus* | 0.6385 | 0.4040 | 0.2345 | **0.0027** | 0.5393 | 0.0993 | 0.0810 |
| Collared dove, *Streptopelia decaocto* | 0.5525 | 0.5649 | -0.0124 | 0.8790 | 0.3876 | 0.1648 | **0.0092** |
| Starling, *Sturnus spp* | 0.3426 | 0.2574 | 0.0852 | 0.1510 | 0.3534 | -0.0108 | 0.8140 |
| Eurasian blackbird, *Turdus merula* | 0.4717 | 0.5500 | -0.0783 | 0.3100 | 0.3683 | 0.1034 | **0.0394** |
| *mean* | *0.5134* | *0.4728* | *0.0406* |  | *0.3864* | *0.1270* |  |
| *SD* | *0.1789* | *0.1508* | *0.1182* |  | *0.1167* | *0.1830* |  |

**Table S4.** *P*-values for the effects of hour, the interaction between hour and group, and the interaction between time and group in the occupancy models. *f* refers to the basis function used to model the non-lineal effect of hour on bird detectability.

| Species | *f*(hour) | *f*(hour)*group | time*group |
| --- | --- | --- | --- |
| European goldfinch, *Carduelis carduelis* | <0.00001 | 0.14308 | 0.04767 |
| Greenfinch, *Carduelis chloris* | <0.00001 | 0.00291 | 0.02069 |
| Rock pigeon, *Columba livia* | 0.00754 | 0.00137 | 0.04995 |
| Wood pigeon, *Columba palumbus* | <0.00001 | 0.00001 | 0.93144 |
| Blue tit, *Cyanistes caeruleus* | <0.00001 | 0.00002 | 0.94749 |
| Yellow-legged gull, *Larus michahellis* | 0.00256 | 0.00001 | 0.00328 |
| White wagtail, *Motacilla alba* | 0.09982 | 0.00018 | 0.00001 |
| Monk parakeet, *Myiopsitta monachus* | 0.00455 | 0.00008 | 0.83579 |
| Great tit, *Parus major* | <0.00001 | 0.00113 | 0.00023 |
| House sparrow, *Passer domesticus* | <0.00001 | 0.00073 | 0.64000 |
| Black redstart, *Phoenicurus ochruros* | 0.01681 | 0.11286 | 0.05613 |
| Magpie, *Pica pica* | 0.00048 | 0.00009 | 0.02690 |
| Serin, *Serinus serinus* | <0.00001 | 0.00027 | 0.01794 |
| Collared dove, *Streptopelia decaocto* | 0.00003 | 0.00039 | 0.02776 |
| Starling, *Sturnus spp* | <0.00001 | 0.00594 | 0.04043 |
| Eurasian blackbird, *Turdus merula* | <0.00001 | 0.00464 | 0.00649 |

**Table S5.** Results for the effects of the interaction between time and group on the detection part of the occupancy models. Columns lockdown, urban and non-urban show the probability of detection of the species after 1 hour of survey. A probability of 0.5 means that the observers may or may not detect the species with the same probability (i.e. flat slope). Except the rock pigeon in urban checklists, in all cases, as expected, these probabilities were above 0.5 (i.e., increased probability of detection of a species throughout sampling time). For the lockdown estimates, the *p*-value testing whether or not this slope was different from 0.5 is shown. However, in urban and non-urban groups, the *p*-value shows whether or not these slopes differed from the lockdown group. The difference (Δ) between these slopes is also provided to enhance result interpretation.

| Species | Lockdown<br>(2020) | $p$ | Urban<br>(2015-19) | $\Delta$ lockdown-<br>urban | $p$ | Non-urban<br>(2015-19) | $\Delta$ lockdown-<br>non-urban | $p$ |
| --- | --- | --- | --- | --- | --- | --- | --- | --- |
| European goldfinch, *Carduelis carduelis* | 0.5372 | 0.1908 | 0.6620 | -0.1248 | **0.0019** | 0.6345 | -0.0973 | **0.0012** |
| Greenfinch, *Carduelis chloris* | 0.5889 | **0.0023** | 0.7675 | -0.1786 | **0.0002** | 0.6282 | -0.0394 | 0.2063 |
| Rock pigeon, *Columba livia* | 0.5250 | 0.6086 | 0.4735 | 0.0515 | 0.4093 | 0.6112 | -0.0862 | 0.0861 |
| Wood pigeon, *Columba palumbus* | 0.6805 | **<0.0001** | 0.6757 | 0.0048 | 0.9130 | 0.6920 | -0.0115 | 0.7310 |
| Blue tit, *Cyanistes caeruleus* | 0.6710 | **<0.0001** | 0.6800 | -0.0091 | 0.8400 | 0.6863 | -0.0154 | 0.6480 |
| Yellow-legged gull, *Larus michahellis* | 0.6936 | **0.0001** | 0.7120 | -0.0184 | 0.7438 | 0.5537 | 0.1399 | **0.0061** |
| White wagtail, *Motacilla alba* | 0.6839 | **<0.0001** | 0.8566 | -0.1726 | **0.0001** | 0.6107 | 0.0733 | **0.0130** |
| Monk parakeet, *Myiopsitta monachus* | 0.6371 | **0.0008** | 0.6298 | 0.0073 | 0.9110 | 0.6685 | -0.0314 | 0.4990 |
| Great tit, *Parus major* | 0.5682 | **0.0197** | 0.7988 | -0.2306 | **<0.0001** | 0.7243 | -0.1560 | **<0.0001** |
| House sparrow, *Passer domesticus* | 0.6983 | **0.0001** | 0.6602 | 0.0381 | 0.5410 | 0.6384 | 0.0599 | 0.2120 |
| Black redstart, *Phoenicurus ochruros* | 0.5089 | 0.7519 | 0.6446 | -0.1357 | **0.0019** | 0.5965 | -0.0876 | **0.0041** |
| Magpie, *Pica pica* | 0.5584 | 0.0532 | 0.7258 | -0.1674 | **0.0005** | 0.6599 | -0.1014 | **0.0021** |
| Serin, *Serinus serinus* | 0.6862 | **<0.0001** | 0.8111 | -0.1248 | **0.0114** | 0.6536 | 0.0327 | 0.4020 |
| Collared dove, *Streptopelia decaocto* | 0.7411 | **<0.0001** | 0.6514 | 0.0897 | 0.1240 | 0.5881 | 0.1530 | **0.0006** |
| Starling, *Sturnus spp* | 0.7244 | **<0.0001** | 0.6823 | 0.0421 | 0.3050 | 0.6212 | 0.1032 | **0.0013** |
| Eurasian blackbird, *Turdus merula* | 0.5705 | **0.0234** | 0.7883 | -0.2178 | **<0.0001** | 0.6726 | -0.1021 | **0.0018** |

## References

1. Seto KC, Güneralp B, Hutyra LR. 2012 Global forecasts of urban expansion to 2030 and direct impacts on biodiversity and carbon pools. Proc. Natl. Acad. Sci. U.S.A. 109,16083–16088. (doi:10.1073/pnas.1211658109)

2. Grimm NB, et al. 2008 Global change and the ecology of cities. Science 319,756– 760. (doi:10.1126/science.1150195)

3. Pickett ST, et al. 2011 Urban ecological systems: Scientific foundations and a decade of progress. J. Environ. Manage. 92,331–362. (doi:10.1016/j.jenvman.2010.08.022)

4. McDonnell MJ, Hahs AK. 2015 Adaptation and adaptedness of organisms to urban environments. Annu. Rev. Ecol. Evol. S. 46,261–280. (doi:10.1146/annurev-ecolsys-112414-054258)

5. Sol D, Lapiedra O, González-Lagos C. 2013 Behavioural adjustments for a life in the city. Anim. Behav. 85,1101–1112. (doi:10.1016/j.anbehav.2013.01.023)

6. Isaksson C. 2018 Impact of urbanization on birds. In Bird species. How they arise, modify and vanish (ed. DT Tietze), pp 235–257. Cham, Switzerland: Springer. (doi:10.1007/978-3-319-91689-7_13)

7. Murgui E, Hedblom M. 2017 Ecology and conservation of birds in urban environments. Cham, Switzerland: Springer. (doi: 10.1007/978-3-319-43314-1)

8. Slabbekoorn H, Ripmeester EA. 2007 Birdsong and anthropogenic noise: implications and applications for conservation. Mol. Ecol. 17, 72–83. (doi:10.1111/j.1365-294X.2007.03487.x)

9. Ortega CP. 2012 Effects of noise pollution on birds: A brief review of our knowledge. Ornithol. Monogr. 74,6–22. (doi:10.1525/om.2012.74.1.6)

10. Rich C, Longcore T. 2006 Ecological consequences of artificial night lighting. Washington, DC: Island Press.

11. Gaston KJ, Bennie J, Davies TW, Hopkins J. 2013 The ecological impacts of nighttime light pollution: a mechanistic appraisal. Biol. Rev. 88, 912–927. (doi:10.1111/brv.12036)

12. Lowry H, Lill A, Wong BB. 2013 Behavioural responses of wildlife to urban environments. Biol. Rev. 88,537–549. (doi:10.1111/brv.12012)

13. Bradley CA, Altizer S. 2007 Urbanization and the ecology of wildlife diseases. Trends Ecol. Evol. 22,95–102. (doi:10.1016/j.tree.2006.11.001)

14. Jiménez-Peñuela J, Ferraguti M, Martínez-de la Puente J, Soriguer R, Figuerola J. 2019 Urbanization and blood parasite infections affect the body condition of wild birds. Sci. Total Environ. 651, 3015–3022. (doi:10.1016/j.scitotenv.2018.10.203)

15. Loss SR, Will T, Marra PP. 2015 Direct mortality of birds from anthropogenic causes. Annu. Rev. Ecol. Evol. S. 46,99–120. (doi:10.1146/annurev-ecolsys-112414-054133)

16. Pavisse R, Vangeluwe D, Clergeau P. 2019 Domestic cat predation on garden birds: An analysis from European ringing programmes. Ardea 107,103–109. (doi:10.5253/arde.v107i1.a6)

17. Aymí R, González Y, López T, Gordo O. 2017 Bird-window collisions in a city on the Iberian Mediterranean coast during autumn migration. Rev. Cat. Ornitol. 33, 17–28.

18. Johnson MTJ, Munshi-South J. 2017 Evolution of life in urban environments. Science 358,eaam8327 (doi: 10.1126/science.aam8327)

19. Szulkin M, Munshi-South J, Charmantier A. 2020 Urban evolutionary biology. New York, USA: Oxford University Press.

20. Anderson RM, Heesterbeek H, Klinkenberg D, Hollingsworth TD. 2020 How will country-based mitigation measures influence the course of the COVID-19 epidemic? Lancet 395,931–934. (doi:10.1016/S0140-6736(20)30567-5)

21. Gatto M, et al. 2020 Spread and dynamics of the COVID-19 epidemic in Italy: Effects of emergency containment measures. Proc. Natl. Acad. Sci. U.S.A. 117,10484–10491. (doi:10.1073/pnas.2004978117)

22. Aleta A, Moreno Y. 2020 Evaluation of the potential incidence of COVID-19 and effectiveness of containment measures in Spain: a data-driven approach. BMC Med. 18, 157. (doi:10.1186/s12916-020-01619-5)

23. Kraemer MU, et al. 2020 The effect of human mobility and control measures on the COVID-19 epidemic in China. Science 368,493–497. (doi:10.1126/science.abb4218)

24. Rutz C, et al. 2020 COVID-19 lockdown allows researchers to quantify the effects of human activity on wildlife. Nat. Ecol. Evol. 4, 1156–1159. (doi:10.1038/s41559-020-1237-z)

25. Venter ZS, Aunan K, Chowdhury S, Lelieveld J. 2020 COVID-19 lockdowns cause global air pollution declines. Proc. Natl. Acad. Sci. U.S.A. 117, 18984–18990. (doi:10.1073/pnas.2006853117)

26. Briz-Redón Á, Belenguer-Sapiña C, Serrano-Aroca Á. 2021 Changes in air pollution during COVID-19 lockdown in Spain: a multi-city study. J. Environ. Sci. 101, 16–26. (doi:10.1016/j.jes.2020.07.029)

27. Aloi A, et al. 2020 Effects of the COVID-19 lockdown on urban mobility: Empirical evidence from the city of Santander (Spain). Sustainability 12,3870. (doi:10.3390/su12093870)

28. Lecocq T, et al. 2020 Global quieting of high-frequency seismic noise due to COVID-19 pandemic lockdown measures. Science 369,1338–1343. (doi:10.1126/science.abd2438)

29. Aletta F, Oberman T, Mitchell A, Tong H, Kang J. 2020 Assessing the changing urban sound environment during the COVID-19 lockdown period using short-term acoustic measurements. Noise Mapping 7,123–134. (doi:10.1515/noise-2020-0011)

30. Asensio C, Pavón I, De Arcas G. 2020 Changes in noise levels in the city of Madrid during COVID-19 lockdown in 2020. J. Acoust. Soc. Am. 148,1748–1755. (doi: 10.1121/10.0002008)

31. Corlett RT, et al. 2020 Impacts of the coronavirus pandemic on biodiversity conservation. Biol. Conserv. 246,108571. (doi:10.1016/j.biocon.2020.108571)

32. Bates AE, Primack RB, Moraga P, Duarte CM. 2020 COVID-19 pandemic and associated lockdown as a “Global Human Confinement Experiment” to investigate biodiversity conservation. Biol. Conserv. 248, 108665. (doi:10.1016/j.biocon.2020.108665)

33. Zellmer AJ et al. 2020 What can we learn from wildlife sightings during the COVID-19 global shutdown?. Ecosphere 11,e03215. (doi:10.1002/ecs2.3215)

34. Manenti R, et al. 2020 The good, the bad and the ugly of COVID-19 lockdown effects on wildlife conservation: Insights from the first European locked down country. Biol. Conserv. 249, 108728. (doi:10.1016/j.biocon.2020.108728)

35. Montgomery RA, Raupp J, Parkhurst M. 2021 Animal behavioral responses to the COVID-19 quietus. Trends Ecol. Evol. 36,184–186. (doi:10.1016/j.tree.2020.12.008)

36. Orro A, Novales M, Monteagudo Á, Pérez-López JB, Bugarín MR. 2020 Impact on city bus transit services of the COVID–19 lockdown and return to the new Normal: The case of A Coruña (Spain). Sustainability 12,7206. (doi:10.3390/su12177206)

37. Saladié Ò, Bustamante E, Gutiérrez A. 2020 COVID-19 lockdown and reduction of traffic accidents in Tarragona province, Spain. TRIP 8,100218. (doi: 10.1016/j.trip.2020.100218)

38. Vardi R, Berger-Tal O, Roll U. 2021 iNaturalist insights illuminate COVID-19 effects on large mammals in urban centers. Biol. Conserv. 254,108953. (doi:10.1016/j.biocon.2021.108953)

39. Simons TR, Alldredge MW, Pollock KH, Wettroth JM. 2007 Experimental analysis of the auditory detection process on avian point counts. Auk 124,986–999. (doi:10.1093/auk/124.3.986)

40. Pacifici K, Simons TR, Pollock KH. 2008 Effects of vegetation and background noise on the detection process in auditory avian point-count surveys. Auk 125,600–607. (doi:10.1525/auk.2008.07078)

41. Díaz M, Parra A, Gallardo C. 2011 Serins respond to anthropogenic noise by increasing vocal activity. Behav. Ecol. 22, 332–336. (doi:10.1093/beheco/arq210)

42. Gil D, Honarmand M, Pascual J, Pérez-Mena E, Macías García C. 2015 Birds living near airports advance their dawn chorus and reduce overlap with aircraft noise. Behav. Ecol. 26, 435–443. (doi:10.1093/beheco/aru207)

43. Rose S, Suri J, Brooks M, Ryan PG. 2020 COVID-19 and citizen science: lessons learned from southern Africa. Ostrich 91,188–191. (doi:10.2989/00306525.2020.1783589)

44. Randler C, Tryjanowski P, Jokimäki J, Kaisanlahti-Jokimäki ML, Staller N. 2020 SARS-CoV2 (COVID-19) Pandemic lockdown influences nature-based recreational activity: The case of birders. Int. J. Environ. Res. Public Health 17,7310. (doi:10.3390/ijerph17197310)

45. Hochachka WM, Alonso H, Gutiérrez-Expósito C, Miller E, Johnston A. 2021 Regional variation in the impacts of the COVID-19 pandemic on the quantity and quality of data collected by the project eBird. Biol. Conserv. 254, 108974. (doi:10.1016/j.biocon.2021.108974)

46. González-Guerrero O, Pons X. 2020 The 2017 Land Use/Land Cover Map of Catalonia based on Sentinel-2 images and auxiliary data. Spanish J. Remote Sens. 55, 81–92. (doi:10.4995/raet.2020.13112)

47. R Core Team. 2017 R: A Language and Environment for Statistical Computing (version 3.4.3). Vienna, Austria: R Foundation for Statistical Computing. (https://www.Rproject.org/)

48. Herrando S, Weiserbs A, Quesada J, Ferrer X, Paquet JI. 2012 Development of an urban bird indicator: using data from monitoring schemes in two large European cities. Anim. Biodivers. Conserv. 35,141–150.

49. Estrada J, Pedrocchi V, Brotons L, Herrando S. 2004 Atles dels ocells nidificants de Catalunya 1999-2002. Barcelona, Spain: ICO-Lynx.

50. Shirihai H, Svensson L. 2018 Handbook of Western Palearctic Birds, Volume II: Flycatchers to Buntings. London, UK: Helm.

51. MacKenzie DI, Nichols JD, Royle JA, Pollock KH, Bailey LL, Hines JA. 2006 Occupancy estimation and modeling: Inferring patterns and dynamics of species occurrence. Amsterdam: Elsevier.

52. Altwegg R, Nichols JD. 2019 Occupancy models for citizen-science data. Methods Ecol. Evol. 10, 8–21. (doi 10.1111/2041-210X.13090)

53. Schmidt JH, McIntyre CL, MacCluskie MC. 2013 Accounting for incomplete detection: What are we estimating and how might it affect long-term passerine monitoring programs? Biol. Conserv. 160, 130–139. (doi:10.1016/j.biocon.2013.01.007)

54. Johnston A, Fink D, Hochachka WM, Kelling S. 2018 Estimates of observer expertise improve species distributions from citizen science data. Methods Ecol. Evol. 9, 88–97 (doi:10.1111/2041-210X.12838)

55. Wood SN. 2017 Generalized Additive Models: An Introduction with R. Boca Raton, FL: CRC Press.

56. Fiske I, Chandler R. 2011 unmarked: An R package for fitting hierarchical models of wildlife occurrence and abundance. J. Stat. Softw. 43, 1–23. (doi:10.18637/jss.v043.i10)

57. Senar JC, Domènech J, Arroyo L, Torre I, Gordo O. 2016 An evaluation of monk parakeet damage to crops in the metropolitan area of Barcelona. Anim. Biodivers. Conserv. 39, 141–145. (doi: 10.32800/abc.2016.39.0141)

58. Covas L, Senar JC, Roqué L, Quesada J. 2017 Records of fatal attacks by Rose-ringed parakeets”Psittacula krameri” on native avifauna. Rev. Cat. Ornitol. 33: 45– 49.

59. Senar JC, Navalpotro H, Pascual J, Montalvo T. 2021 Nicarbazin has no effect on reducing feral pigeon populations in Barcelona. Pest Manag. Sci. 77: 131–137. (https://doi.org/10.1002/ps.6000)

60. Pocino N, Giralt N, Ferrer X. 2005 Colonization and expansion of the Collared Dove Streptopelia decaocto in Catalonia. Rev. Cat. Ornitol. 21: 1–10.

61. ICO. 2020 SIOC: servidor d’informació ornitològica de Catalunya. Barcelona: ICO. (http://www.sioc.cat)

62. Nordt A, Klenke R. 2013 Sleepless in town–drivers of the temporal shift in dawn song in urban European blackbirds. PloS ONE 8,e71476. (doi:10.1371/journal.pone.0071476)

63. Sierro J, Schloesing E, Pavón I, Gil D. 2017 European blackbirds exposed to aircraft noise advance their chorus, modify their song and spend more time singing. Front. Ecol. Evol. 5, 68 (doi:10.3389/fevo.2017.00068)

64. Derryberry EP, Phillips JN, Derryberry GE, Blum MJ, Luther D. 2020 Singing in a silent spring: Birds respond to a half-century soundscape reversion during the COVID-19 shutdown. Science 370,575–579. (doi:10.1126/science.abd5777)

65. Gilby BL, et al. 2021 Potentially negative ecological consequences of animal redistribution on beaches during COVID-19 lockdown. Biol. Conserv. 253, 108926. (doi:10.1016/j.biocon.2020.108926)

66. Soh MC, Pang RY, Ng BX, Lee BPH, Loo AH, Kenneth BH. 2021 Restricted human activities shift the foraging strategies of feral pigeons (Columba livia) and three other commensal bird species. Biol. Conserv. 253, 108927. (doi:10.1016/j.biocon.2020.108927)

67. Hentati-Sundberg J, Berglund PA, Hejdström A, Olsson O. 2021 COVID-19 lockdown reveals tourists as seabird guardians. Biol. Conserv. 254, 108950. (doi:10.1016/j.biocon.2021.108950)

68. Frédéric, L., et al. 2021 COVID19-induced reduction in human disturbance enhances fattening of an overabundant goose species. Biol. Conserv. 255, 108968. (doi:10.1016/j.biocon.2021.108968)

69. Gordo O, Sanz JJ, Lobo JM. 2008 Geographic variation in onset of singing among populations of two migratory birds. Acta Oecol. 34, 50–64. (doi:10.1016/j.actao.2008.03.006)

70. Richards DG. 1981 Environmental acoustics and censuses of singing birds. Stud. Avian Biol. 6, 297–300.

71. Brown TJ, Handford P. 2003 Why birds sing at dawn: the role of consistent song transmission. Ibis 145,120–129. (doi:10.1046/j.1474-919X.2003.00130.x)

72. Arroyo-Solís A, Castillo JM, Figueroa E, López-Sánchez JL, Slabbekoorn H. 2013 Experimental evidence for an impact of anthropogenic noise on dawn chorus timing in urban birds. J. Avian Biol. 44,288–296. (doi:10.1111/j.1600-048X.2012.05796.x)

73. Cartwright LA, Taylor DR, Wilson DR, Chow-Fraser P. 2014 Urban noise affects song structure and daily patterns of song production in Red-winged Blackbirds (Agelaius phoeniceus). Urban Ecosyst. 17, 561–572. (doi:10.1007/s11252-013-0318-z)

74. Fuller RA, Warren PH, Gaston KJ. 2007 Daytime noise predicts nocturnal singing in urban robins. Biol. Lett. 3, 368–370. (doi:10.1098/rsbl.2007.0134)

75. Planesas P. 2013 La hora oficial en España y sus cambios. Anuario del Observatorio Astronómico de Madrid 1,373–404.

76. Shannon G, et al. A synthesis of two decades of research documenting the effects of noise on wildlife. Biol. Rev. 91, 982–1005. (doi:10.1111/brv.12207)

77. Houston AI, McNamara JM. 1993 A theoretical investigation of the fat reserves and mortality levels of small birds in winter. Ornis Scandinavica 24,205–219. (doi:10.2307/3676736)

78. Villén-Pérez S, Carrascal LM, Gordo O. 2014 Wintering forest birds roost in areas of higher sun radiation. Eur. J. Wildlife Res. 60, 59–67. (doi:10.1007/s10344-013-0750-7)

79. Bautista LM et al. 2004 Effect of weekend road traffic on the use of space by raptors. Conserv. Biol. 18, 726–732. (doi:10.1111/j.1523-1739.2004.00499.x)

80. Guillera-Arroita G, Lahoz-Monfort JJ, MacKenzie DI, Wintle BA, McCarthy MA. 2014 Ignoring imperfect detection in biological surveys is dangerous: A response to ‘Fitting and interpreting occupancy models’. PLoS ONE 9,e99571. (doi:10.1371/journal.pone.0099571)

81. Herrando S, Estrada J, Brotons L, Guallar S. 2016 Do common bird winter censuses produce similar results when conducted in the morning and in the afternoon? Rev. Cat. Ornitol. 22, 14–20.

